# Binding and unbinding pathways of peptide substrate on SARS-CoV-2 3CL protease

**DOI:** 10.1101/2022.06.08.495396

**Authors:** Kei Moritsugu, Toru Ekimoto, Mitsunori Ikeguchi, Akinori Kidera

**Affiliations:** Graduate School of Medical Life Science, Yokohama City University, 1-7-29 Suehirocho, Tsurumi, Yokohama 230-0045, Japan; Graduate School of Science, Osaka Metropolitan University, 1-2 Gakuencho, Naka-ku, Sakai, Osaka 599-8570, Japan

## Abstract

Based on many crystal structures of ligand complexes, much study has been devoted to understanding the molecular recognition of SARS-CoV-2 3C-like protease (3CL^pro^), a potent drug target for COVID-19. In this research, to extend this present static view, we examined the kinetic process of binding/unbinding of an eight-residue substrate peptide to/from 3CL^pro^ by evaluating the path ensemble with the weighted ensemble simulation. The path ensemble showed the mechanism of how a highly flexible peptide folded into the bound form. At the early stage, the dominant motion was the diffusion on the protein surface showing a broad distribution, whose center was led into the cleft of the Chymotrypsin fold. We observed a definite sequential formation of the hydrogen bonds at the later stage occurring in the cleft, initiated between Glu166 (3CL^pro^) and P3_Val (peptide), followed by binding to the oxyanion hole and completed by the sequencespecific recognition at P1_Gln.

## Introduction

The 3C-like protease (3CL^pro^, also called the main protease) of SARS-CoV-2 is a cysteine protease of the C30 family (1) and it plays a critical role in viral replication by cleaving the polyprotein of SARS-CoV-2 at 11 distinct sites to release functional proteins (Fig. 1a; 2). Thus, 3CL^pro^ is a potent drug target that has been widely investigated for preventing the COVID-19 pandemic (2–4). A number of crystal structures complexed with the substrate peptides have been solved for the atomic-level understanding of the proteolytic mechanism of 3CL^pro^ (5–12) and it has been shown that the substrate recognition of 3CL^pro^ can be outlined by the classic subsite model of proteases (13,14). This knowledge has been the basis for the inhibitor design (3,4), particularly for the peptidomimetic inhibitors (15,16).

**Fig. 1.**
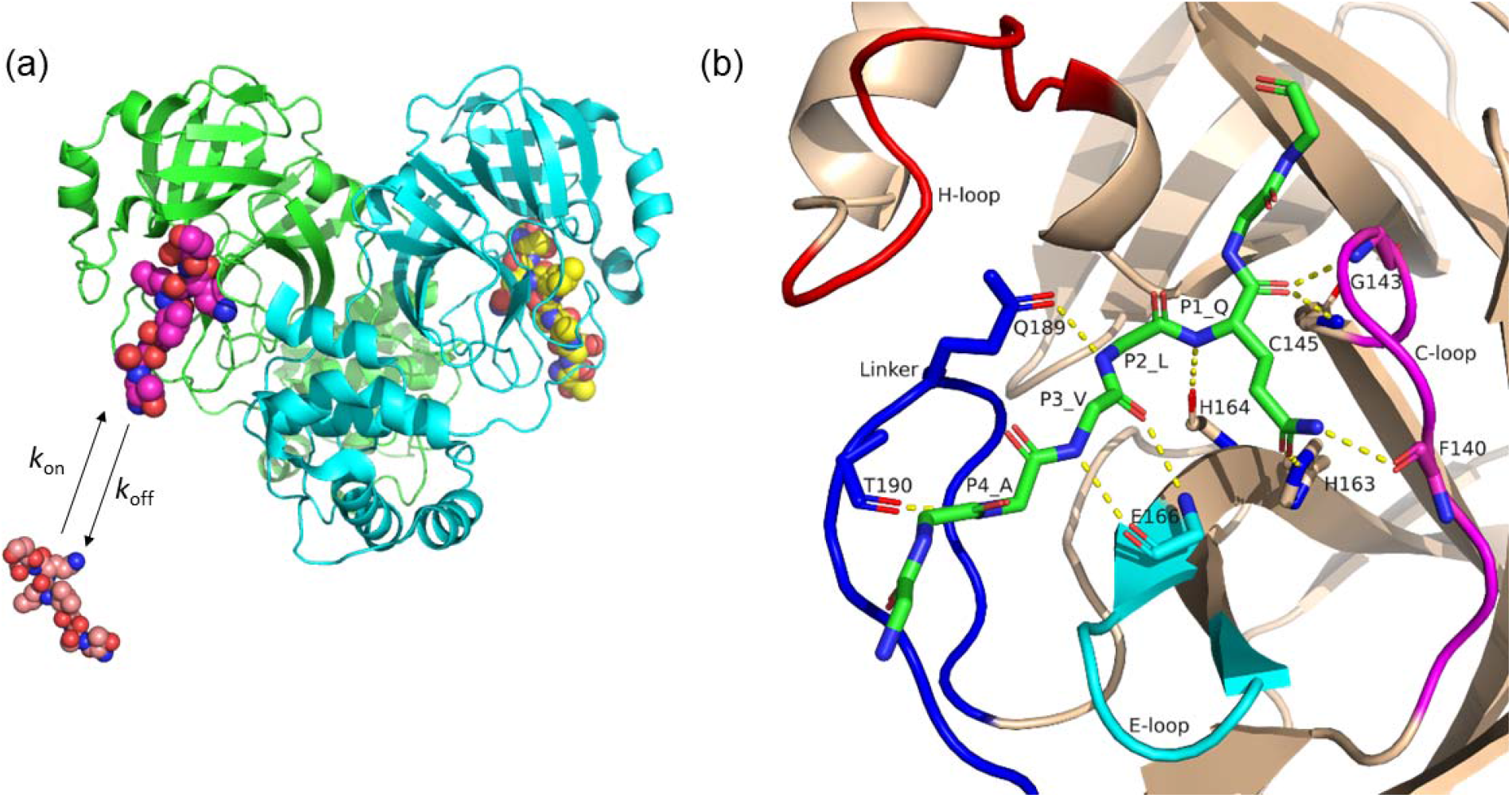
SARS-CoV-2 3CL^pro^ structure. (a) The 3CL^pro^ dimeric structure in complex with the eight-residue peptide (sphere) bound on each protomer (PDB: 2q6g, 34), where the peptide’s three C-terminal residues are excluded in the simulation model. The peptide is coloured magenta (bound on protomer A, by green) and yellow (bound on protomer B, by cyan). The peptide binding/unbinding process to/from protomer A (*k*_on_ and *k*_off_ are the associated rate constants, respectively) were simulated while protomer B was kept in the peptide-bound state. (b) A closeup view indicates hydrogen bonds between 3CL^pro^ and the peptide (Table 1) by dashed yellow lines together with the peptide by stick representation. The moving clusters defined in the Motion Tree (see Figs. S1 and S2) are also shown: H-loop (43–51: red), C-loop (138–144: magenta), E-loop (166–171: cyan) and Linker (184–197: blue). Domains 1 and 2 are in wheat and grey, respectively. The peptide is bound at the cleft between the two domains.

In our previous study (17), the structural changes of 3CL^pro^ upon ligand binding were discussed collectively by accumulating all available crystal structures of 3CL^pro^ including both ligand-bound and ligand-free forms (more than 300 PDB entries, including homologous 3CL^pro^ of SARS-CoV, comprising a broad range of the conformations). We call the assembled crystal structures ‘*crystal structure ensemble*’ (17,18). It was found in the “*crystal structure ensemble*’ that the structural responses upon ligand binding exclusively occurred at the ligand binding flexible loops on the rigid Chymotrypsin fold to finely regulate the catalytic activity (Fig. 1b; 17).

In this research, we extend the scope of the study on 3CL^pro^ from thermodynamics to kinetics to obtain the atomic details of the binding/unbinding process by employing the path sampling simulations (19–21). The kinetic view offers a significantly larger amount of information describing the entire process of the molecular recognition compared to the thermodynamic view focusing only on the initial (ligand-free) state and the final (ligand-bound) state. Recently, the kinetic information has attracted considerable attention because the drug efficacy *in vivo* was found to be more relevant to the kinetics rather than the thermodynamics in the equilibrium condition (22,23).

The kinetics of the peptide binding process is sensitively modulated by different factors, not only the peptide’s fluctuations but also the protein’s structural changes and desolvation in the interface. Then, the path sampling of binding and unbinding processes must be conducted in a sufficiently comprehensive manner to produce a trajectory covering all events occurring in all degrees of freedom, including those of solvent molecules. Since such a molecular simulation is computationally highly demanding, we used the weighted ensemble (WE) simulation (24–26), which runs a number of short-time unconstrained all-atom MD simulations to evolve a diverse set of continuous paths of the protein structural change efficiently. The path ensemble of the binding/unbinding process thus obtained offers the essential information of the molecular recognition to show the atomistic details of the binding/unbinding process.

We chose the eight-residue peptide (TSAVLQ↓SG, ↓ indicating the cleavage site, see Methods for details) from the 11 cleavage sites of SARS-CoV-2 (2) for the target substrate in the simulations. This is a model for the native substrate as well as for the flexible peptidomimetic compounds (mostly Mw > 500) that are found in the majority of the ligand molecules in the complex crystal structures (17). This computer simulation study elucidates how 3CL^pro^ induces the folding and binding of the highly flexible eight-residue peptide from the fully random conformation in the bulk solvent to the peptide-bound structure uniquely determined at the 3CL^pro^ recognition site, and answers a question as to how the structural changes in the flexible loops of 3CL^pro^ contribute to the peptide recognition process.

## Results

### Structural dynamics of 3CL^pro^ in peptide-bound and peptide-free forms

Prior to the WE simulation, conventional MD simulations of the SARS-CoV-2 3CL^pro^ dimer were conducted with the peptide-bound and peptide-free forms to describe the structural dynamics characteristics for the two terminal states of the path ensemble. This procedure aims at finding the definite structural changes upon ligand binding and identifying the native atom contacts with the peptide under the thermal fluctuation, that are useful for the subsequent analysis of the path ensemble.

To detect the structural difference seen in the two MD simulations, the Motion Tree was constructed from the difference between the two snapshot structures near each average structure obtained from the trajectories of the peptide-bound and peptide-free simulations (27; Fig. S1). The Motion Tree reveals the structural differences between the two termini of the path ensemble that mostly occurs in the ‘moving clusters’ (the segments showing large structural changes) originally defined by the crystal structure ensemble (17), i.e. ‘H-loop’ (residues 38–64), ‘E-loop’ (residues 166–178), ‘Linker’ (residues 188–196) and ‘domain III’ (residues 200–300) (these clusters were named the same as in ref. 17). ‘C-loop’ (residues 138–143) was another major moving cluster observed in the crystal structure ensemble, while in Fig. S1, it is not a major cluster but is found in a part of the rigid core region with a small MT score less than 1 Å. This is because the crystal structure ensemble contains both the active and collapsed (inactive) structures of the C-loop, whose pairwise Cα RMSD exceeds 3Å, corresponding to the MT score between nodes 3 and 4 in Fig. 2. However, the present simulations started with the same crystal structure in the active state (PDB: 2q6g) and the active structure remained stable in the simulation condition even when the peptide was removed. The C-loop’s stability can also be confirmed by the five hydrogen bonds (HBs) between the C-loop and other parts of the protein, found in the crystal structures (Table S1; in the crystal structures, these HBs were formed in the active state but not in the collapsed state; 17); the C-loop will be intrinsically in active form because collapsed C-loop is found only in the crystal structures with mutations at the important sites or appended amino acids at the N-terminus (17). The HBs in the simulations are independent of peptide binding, seen in almost the same probability of occurrence (the average probabilities are 0.74 and 0.77 for the peptide-bound and peptide-free simulations, respectively).

**Fig. 2.**
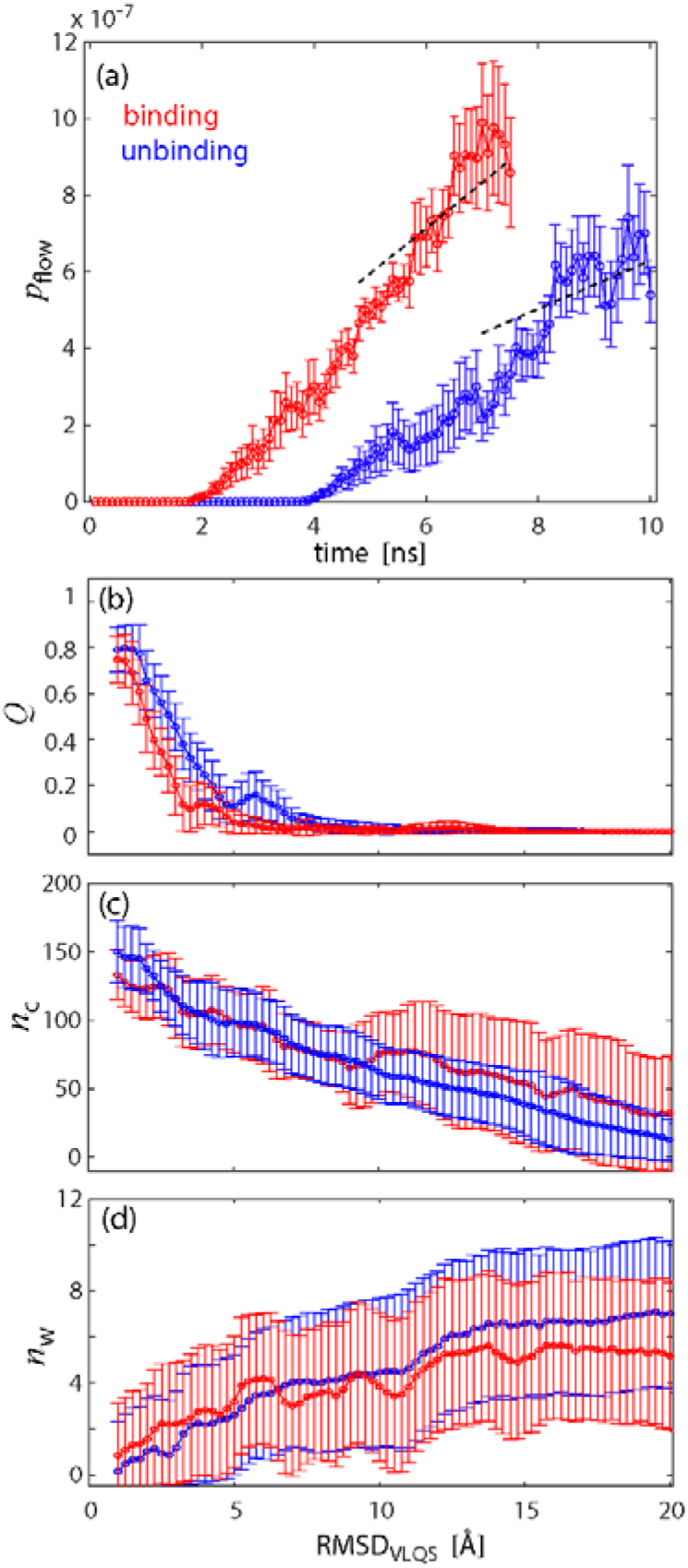
Peptide binding and unbinding WE simulations. (a) Probability flows, *p*_flow_, between the peptide-bound and peptide-free states during the WE simulations of the peptide binding (red) and unbinding (blue). Linear regression lines for the MFPT estimation are shown by dashed lines. The peptide-bound state in this figure was redefined by the conditions that the two HBs in group 2 and more than one HB in group 3 in Table 1 are formed. The error bars were calculated for the three runs of the WE simulations starting at different initial conditions. (b) The ratio of the native contacts formed, *Q*, which was defined by the 43 native atom contacts with a distance less than 4 Å and more than 70% probability of occurrence in the peptide-bound MD simulation. (c) The number of atom contacts with the peptide, *n*_C_, is calculated for the pairs of non-hydrogen atoms with *r* < 4 Å between the protein and the residues VLQS of the peptide. (d) The amount of hydrated water in the residues VLQS of the peptide in the form of the increment to the average number of hydrated water molecules during the peptide-bound simulation, i.e. *n*_W_ – <*n*_W(bound)_>, where <*n*_W(bound)_> = 8.1. Binding (red) and unbinding (blue) processes. The mean and the standard deviation were calculated using all the simulation data.

The dynamic fluctuation of the peptide-bound state and that of the peptide-free state were then compared in the Motion Trees (Fig. S2), which were constructed from the variance of the interresidue distances (28). As shown in Fig. S2, the H-loop, Linker and domain III fluctuate most largely both in the peptide-bound and peptide-free states. Thus, the structural differences between the peptide-bound and peptide-free structures of these clusters illustrated in Fig. S1 were not attributed to the influence of peptide binding, but were rather caused by the large fluctuations. Domain III is distant from the peptide binding sites, and the H-loop and Linker have no substantial interactions with the peptide. However, the E-loop is the cluster that was strongly influenced by peptide binding; the E-loop was stiffened by peptide binding (the MT score decreased from 2 Å to 0.4 Å upon binding; Fig. S2). The E-loop’s role in peptide recognition is discussed below. It was also found in the MD simulations that these moving clusters fluctuated independently of each other as in the crystal structure ensemble (Table S2; 17).

The interactions with the peptide were analyzed by measuring the probability of occurrence of the HBs (Table 1). Most of the HBs found in the simulation were those found in the crystal structure ensemble (17), but the HBs with the Linker, HB #8 and #9, have very low values of the probability of occurrence in the simulation; their probabilities are smaller than 0.1. As shown in Fig. S2, where the Linker maintained a large fluctuation even after peptide binding, the large fluctuation of the Linker in the simulation may prevent the peptide from forming stable HBs, while the crystal structure reflects the one with a low level of fluctuation measured at a low temperature. These HBs are employed below in the description of the binding/unbinding pathway.

**Table 1.** Probability of hydrogen-bond formation *p*_HB_ between 3CL^pro^ and peptide.

| Group<sup>a</sup> | No<sup>b</sup> | 3CL<sup>pro</sup> | Peptide<sup>c</sup> | $p_{\text{HB,crystal}}$<sup>d</sup> | $p_{\text{HB,MD}}$<sup>e</sup> |
| --- | --- | --- | --- | --- | --- |
| 1 | 1 | Glu166N | P3_ValO | 0.77 | 1.00 |
|  | 2 | Glu166O | P3_ValN | 0.74 | 1.00 |
| 2 | 3 | Gly143N | P1_GlnO | 0.56 | 0.98 |
|  | 4 | Cys145N | P1_GlnO | 0.96 | 0.79 |
|  | 5 | His164O | P1_GlnN | 0.87 | 0.97 |
| 3 | 6 | Phe140O | P1_GlnN<sub>ε</sub> | 0.63 | 0.67 |
|  | 7 | His163N<sub>ε</sub> | P1_GlnO<sub>ε</sub> | 0.78 | 0.73 |
| 4 | 8 | Gln189O<sub>ε</sub> | P2_LeuN | 0.55 | 0.09 |
|  | 9 | Thr190O | P4_AlaN | 0.28 | 0.07 |

<sup>a</sup> ‘Group’ is the classification of the HBs occurring sequentially along the pathway (see text).
<sup>b</sup> HBs formed in the crystal structures of the complexes with a peptide or a peptide-mimic compound (17).
<sup>c</sup> The position of the amino acid is for the peptide of TSAVLQ↓SG (‘↓’ indicates the substrate cleavage site).
<sup>d</sup> $p_{\text{HB,crystal}}$ is the probability of occurrence of HBs in the crystal structure ensemble (17).
<sup>e</sup> $p_{\text{HB,MD}}$ for the probability of occurrence of HBs observed in the peptide-bound MD simulations.

### Peptide binding and unbinding kinetics observed in WE simulations

The WE simulations were conducted to obtain the kinetic information and the path ensemble of the peptide binding and unbinding processes. The root-mean-square deviation (RMSD) of the non-hydrogen atoms of the substrate residues VLQS from the crystal structure after superimposing the protein’s core region of the protein, RMSD_VLQS_, was taken as the reaction coordinate (see Methods for details). As summarized in Fig. S3, the unbinding paths were computed starting from the position of the crystal structure (Fig. S3a). The peptide dissociated from the binding site to different directions and finally arrived at the peptide-free state in the bulk solvent region (Fig. S3b). The binding WE runs were conducted to obtain the binding paths reaching the binding site, starting from three distinct peptide-free structures taken from the terminal bin of the unbinding WE simulation (Fig. S3c). Finally, the structures in the peptide-bound state were successfully sampled in the terminal bin of the binding WE runs (Fig.S3d).

The key quantity provided by the WE simulation is the probability flow, *p*_flow_, from the peptide-free state to the peptide-bound state and in the reverse direction. Fig. 2a illustrates the time evolution of *p*_flow_ that allows us to assess the mean first passage time (MFPT) from the approximately linear ranges (5 ≤ *t* ≤ 7.5 ns for the binding simulation and 7.5 ≤ *t* ≤ 10 ns for the unbinding simulation), while latent periods appeared in small *t* (< ~2–4 ns) due to finite numbers of copies in the WE simulation. The kinetic constants, *k*_on_ and *k*_off_, calculated with the estimated peptide concentration, turned out to be *k*_on_ = 6.7 ± 0.71 × 10^4^ M^−1^sec^−1^ and *k*_off_ = 63 ± 9.2 sec^−1^ (Eqs. 1 and 2, see Methods). This result also shows that the binding process started from the three structures (see Fig. S3c) is faster (larger probability flow) than the unbinding process that directly corresponds to the binding affinity, or the dissociation constant: *K*_D_ = *k*_off_ / *k*_on_ ~ 900 μM. Although no experimental *K*_D_ value of the substrate is available, the Michaelis constant *K*_M_ is cited as a reference (Table S3; *K*_M_ is a reasonable reference, as the *k*_cat_ values are small). The experimental *K*_M_ values (5-170 μM) are at least five-fold smaller than the calculated *K*_D_ value. This difference is due to the large *k*_off_ value. Several reasons are supposed, such as the smaller chain length of the substrate adopted in the simulation, and such that the fixed force field did not produce a sufficient affinity in the peptide-bound form. The Michaelis complex related to *K*_M_ is a pre-transition state complex containing the charge and proton transfer between the substrate and the protein, which may yield stronger affinity (29). In a neutron diffraction structure (PDB:7jun; 30), the proton transfers from Sγ in Cys145 to Nε in His41 to produce Cys145^−^ and His41^+^.

To view in details the binding/unbinding kinetics along the reaction coordinate, the probability flow, *p*_flow_, was computed for each bin of the WE simulations, and the MFPT as a function of RMSD_VLQS_ was evaluated similarly as in Fig. 2a. Fig. S4 shows that the evolution curves of MFPT along RMSD_VLQS_ for unbinding and binding processes are reasonably approximated by the power-law type relation as MFPT(*r*) ~ *r^d^* (unbinding) and 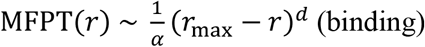, where *r*:= RMSD_VLQS_ with 2.5 Å ≤ *r* ≤ 25.5 Å and *r*_max_ = 28 Å. The power *d* is the effective dimension, which was evaluated here to be ~7 for both processes and α (= 1.5-2.2) is a scaling factor to the absolute values of the MFPT for the two processes.

We can refer to the formulation of the scale-free network (31) for the interpretation of the power-law type behavior, in which the recurrent diffusion occurring in an equilibrium system has the MFPT of the form, MFPT(*r*) ~ *r*^*d*_w_^, where *d*_w_ represents the walk dimension related to the mean square displacement as, 〈Δ*r*^2^〉 ~ *t_β_* with *β* = 2/*d*_w_. According to the formulation, the effective dimension, *d*, evaluated above can be interpreted as the walk dimension *d*_w_, thus *β* = 0.3. Therefore, the dynamics of the binding/unbinding process of the peptide are subdiffusion with *β* < 1. The extensive slowness of the dynamics may come from the flexible torsional motions of the peptide (32) and the interplay with the protein surface restraining the peptide motions.

### Path ensemble of peptide binding and unbinding

Along the binding/unbinding pathways, there are two stages of the process experienced, each of which is described by a different reaction coordinate: the region distant from the binding site, where the peptide moves randomly on the protein surface, is outlined by RMSD_VLQS_ and the region near the binding site, completing the peptide recognition, is explained by the fraction of the native atom contacts between the protein and the peptide, *Q* (see the caption of Fig. 2 for the definition). Fig. 2b illustrates the relation between RMSD_VLQS_ and *Q*. The native contacts occur only in the vicinity of the binding site, RMSD_VLQS_ < 5–7 Å, where *Q* can be employed as the reaction coordinate.

First, the peptide binding and unbinding processes are outlined along the reaction coordinate of RMSD_VLQS_. The structural indices characterising the processes are the number of atom contacts between the protein and the peptide, *n_C_*, and the number of hydrated water molecules of the peptide, *n*_W_ (see the caption of Fig. 2 for the definitions). Fig. 2c illustrates a gradual, almost linear increase in *n*_C_ starting from the non-zero value at RMSD_VLQS_ = 20 Å, showing that at the remote region (RMSD_VLQS_ > 5–7 Å) the peptide has already had plenty of non-specific, nonnative interactions with the protein. This observation indicates that the kinetics in this region is regulated by diffusion on the protein surface. Fig. 2d shows that the peptide is fully solvated in the region of RMSD_VLQS_ > 10 Å. On approaching the peptide-bound state beyond this region, *n*_W_ gradually decreases with RMSD_VLQS_. At the end of this section, it is demonstrated that at RMSD_VLQS_ ~ 10 Å, the binding mode is divided into the inside (< 10 Å) and the outside (> 10 Å) of the cleft of the Chymotrypsin fold in which the binding site exists (see Fig. 1b). Entering the cleft thus accompanies a certain level of desolvation.

As a whole, Figs. 2b-d show that the binding and unbinding processes follow an identical pathway within the margin of error. Thus, we calculated all structural indices without distinguishing the binding and unbinding processes or by mixing up the ensembles of the two processes in the following discussion.

In Fig. 3, we go into detail about the peptide binding process. Fig. 3a illustrates the peptide’s spatial distribution in terms of the translation and rotation degrees of freedom; the vector from the center of Cα atoms of the protein residues, Met165 and Cys145, to the center of Cα atoms of the peptide residues, P2_Val and P0_Ser, is described in the polar coordinates, (*d, θ, φ*). The three-dimensional coordinates are further reduced to the two dimensions, (*d*, |Δ*θ*| + |Δ*φ*|), where Δ shows the difference from the values in the peptide-bound state. The resultant two-dimensional free energy landscape in Fig. 3a shows that when *d* > 10–12 Å, the peptide’s rotation is not hindered by the protein but allowed freely as in the bulk solvent region. When *d* < 10–12 Å, the rotation began to be restricted by the protein surface and was finally reduced to the fluctuation level observed in the peptide-bound simulation. Since *d* is well correlated with RMSD_VLQS_ (see the caption of Fig. 3), the position of *d* ~ 10–12 Å corresponds to RMSD_VLQS_ ~ 10–12 Å where the peptide starts to enter the cleft of the Chymotrypsin fold and its overall rotation starts to be constrained. However, as illustrated in Fig. 3b, even after the peptide enters the cleft, the peptide’s conformation, depicted by Cα-RMSF_int_ (Cα-root-mean-square fluctuation after superimposing the residues VLQS), remains largely fluctuating. It converges to the peptide-bound structure only after the native contacts are formed (RMSD_VLQS_ < 5 Å). It is concluded that the peptide’s conformation is solely determined by the native interactions with the protein. These results are also observed in the peptide’s representative structures (Fig. S5).

**Fig. 3.**
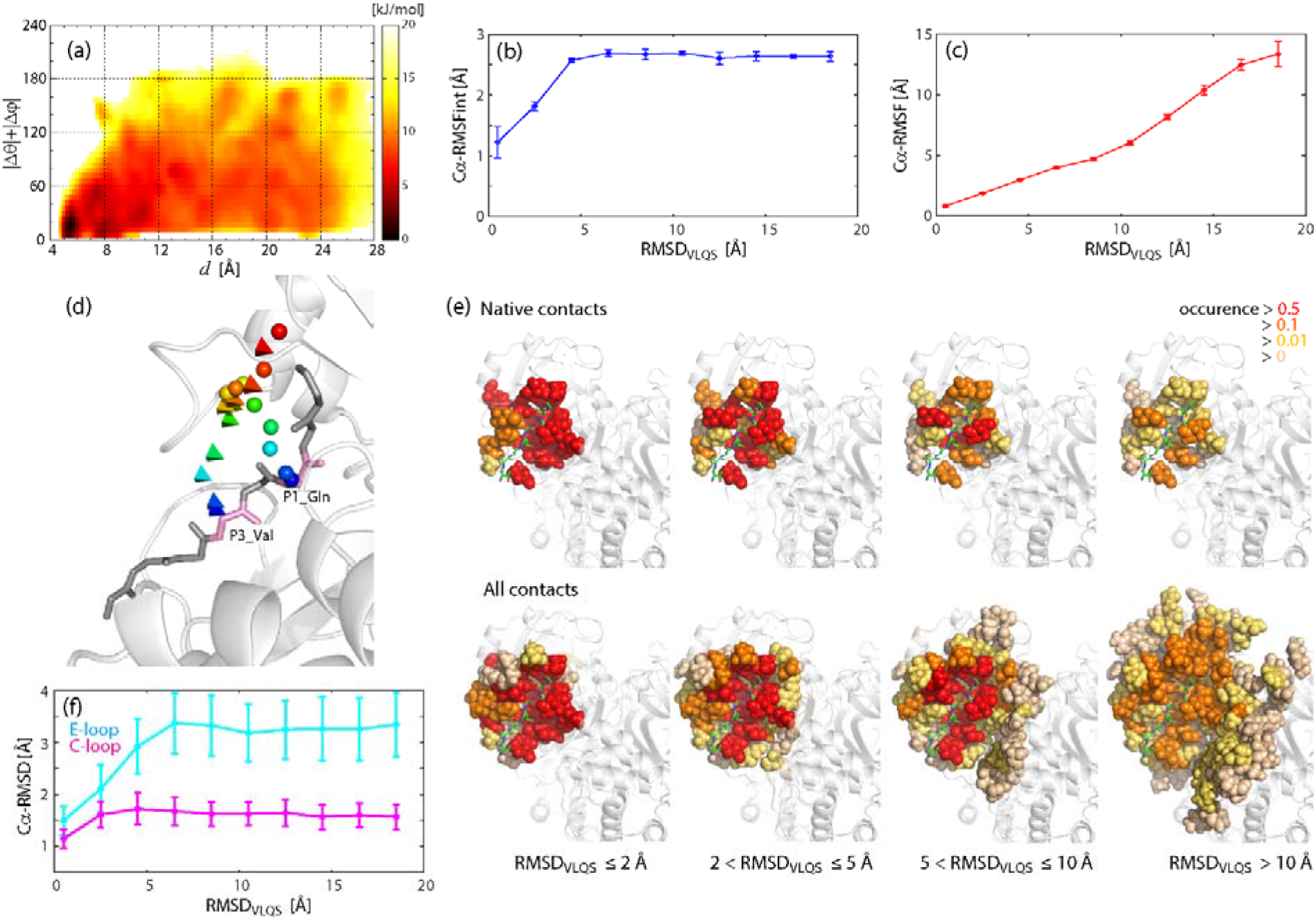
Peptide binding and unbinding processes. (a) Free energy landscape plotted in the plane describing the peptide configuration, represented by the polar coordinates (*d*, *θ,φ*) of the vector connecting the center of Cα of Cys145 and Met165 to the center of Cα of P2_Val and P0_Ser. The horizontal axis *d* is the distance between the peptide and the protein (*d* is related to RMSD_VLQS_ by *d* ≈ (0.93RMSD_VLQS_ + 2) ± 2.5 Å for *d* ≥ 5 Å) and the vertical axis is |Δ*θ*| + |Δ*φ*| representing the rotation of the peptide evaluated by the sum of the absolute deviations of two angles from that of the peptide-bound structure. (b) The peptide’s internal flexibility calculated by the root-mean-square fluctuation (RMSF) of the Cα atoms of the residues VLQS after superimposing VLQS, plotted against RMSD_VLQS_. (c) The width of the distribution of the peptide configuration, calculated by the Cα-RMSF of the peptide residues VLQS after superimposing the core domain as a function of RMSD_VLQS_. In (b) and (c), the error bars were calculated as the standard deviations of the five groups that were generated randomly from the trajectories of the WE simulation. (d) The paths of the Cα atoms on P1_Gln (sphere) and P3_Val (tetrahedra), calculated as their average positions over the simulated peptide structures with a specified range of RMSD_VLQS_. Ten marks are drawn with the colours changing from blue to red, respectively corresponding to the ranges of RMSD_VLQS_ = 0.5 + 2*n* ± 1Å (*n* = 0,…, 9), or 0–1.5 Å (blue), 1.5–3.5 Å,…, 17.5–19.5 Å (red). (e) The 3CL^pro^ residues interacting with the peptide are represented by spheres, separately for each RMSD_VLQS_ range. The colours indicate the ratio of occurrences in the WE simulations. The peptide in the peptide-bound form is shown by green sticks. (f) The Cα-RMSD of the E-loop (cyan) and the C-loop (magenta), plotted against RMSD_VLQS_.

Now let us look into the evolution of the path ensemble from the standpoint of the shift of the peptide distribution. Fig. 3c shows the Cα-RMSF of the residues VLSQ after superimposing the protein’s core region against RMSD_VLQS_. The breadth of the distribution increases with increasing RMSD_VLQS_, with the linear relation, Cα-RMSF ~ 3/4 RMSD_VLQS_. Two extreme scenarios of the binding process can be supposed: (1) the distribution approaching the binding site along a definite pathway and (2) the distribution simply shrinking in size to converge to the binding site without a substantial shift of the distribution center. The first scenario will give a small change in RMSF with RMSD_VLQS_, while the latter gives a substantial change in RMSF. The relation in Fig. 3c may be better suited to the second scenario. This is seen in the distribution of the contact residues in the protein (Fig. 3e), which reveals a shrink of the distribution size along the binding process or with decreasing RMSD_VLQS_. For all ranges of RMSD_VLQS_, the residues surrounding the native binding site are most frequently contacted with the peptide.

It is then possible to scrutinise how the center of the peptide distribution moves with RMSD_VLQS_. Fig. 3d illustrates the evolution, which was monitored separately by the positions of two atoms, Cα atoms of P1_Gln and P3_Val, as a function of RMSD_VLQS_. The center approaches the binding site along the peptide’s longitudinal axis in the peptide-bound state from the P2’ side. It is most notable that the two positions of P1_Gln and P3_Val merge after RMSD_VLQS_ > 10 Å and diverge to each binding position when RMSD_VLQS_ < 10 Å. This indicates that the region of RMSD_VLQS_ ~ 10 Å separates the inside and outside of the cleft of the Chymotrypsin fold, and at the outside of the cleft, the peptide’s overall rotation mixes up the positions of the two Cα atoms. Once the peptide goes into the cleft, the rotation becomes restricted and the two Cα atoms are aligned to the cleft.

Finally, in this section, the ligand-induced structural changes in the protein are discussed. Fig. 3f illustrates the Cα-RMSD value of the E-loop and C-loop. As in the Motion Tree of Fig. S1, the E-loop has a definite structural change, while the C-loop shows only a small change. At RMSD_VLQS_ ~ 6.5 Å, the E-loop’s RMSD value starts to decrease, indicating that the E-loop’s structure is affected mostly by the native contact with the peptide. A more detailed discussion is presented in the next section in terms of the *Q* dependence. However, the H-loop and Linker maintain large fluctuations and exhibit almost no effect on the RMSD_VLQS_ values from peptide binding (Fig. S6a, see also Fig. S2). However, the Linker’s *n*_C_ value increases with a decrease in RMSD_VLQS_ (Fig. S6b). Since the Linker does not have any native contact counted in *Q* (see also Table 1), it was concluded that the large fluctuating Linker makes a substantial number of nonspecific contacts with the peptide (corresponding to 25%–35% of the total number of contacts shown in Fig. 2c), which certainly contributes to not only the binding affinity but also the substrate peptide’s kinetics. The same type of non-specific interaction may occur in the recognition of the peptide-mimic compound with the Linker.

### Completion of the peptide binding to 3CL^pro^

The binding/unbinding process near the binding site was studied in detail along the reaction coordinate of *Q*, the fraction of the native contacts. The structural indices focused here is the peptide recognition by the HBs defined in the peptide-bound simulation (see Fig. 1b and Table 1). In terms of the peptide’s binding sites, the HBs were classified into four groups: (1) P3_Val, (2) P1_Gln (main-chain; the carbonyl oxygen is the target of the oxyanion hole), (3) P1_Gln (side-chain; the origin of the sequence-specific recognition) and (4) P2_Leu and P4_Ala. Group 4 is not discussed here because of the low probability of occurrence in the simulation. The average number of the HBs in each group, *n*_HB_, illustrated in Fig. 4a, indicates that there exists a definite sequence of the formation of the HBs during the peptide binding/unbinding process; the groups 1, 2 and 3 in the ascending order of *Q*. It is again noted that the binding and unbinding processes are mixed for the calculation of *n*_HB_. The difference between the two processes along *Q* is small enough for both processes to keep the sequence of the HB formation (Fig. S7).

**Fig. 4.**
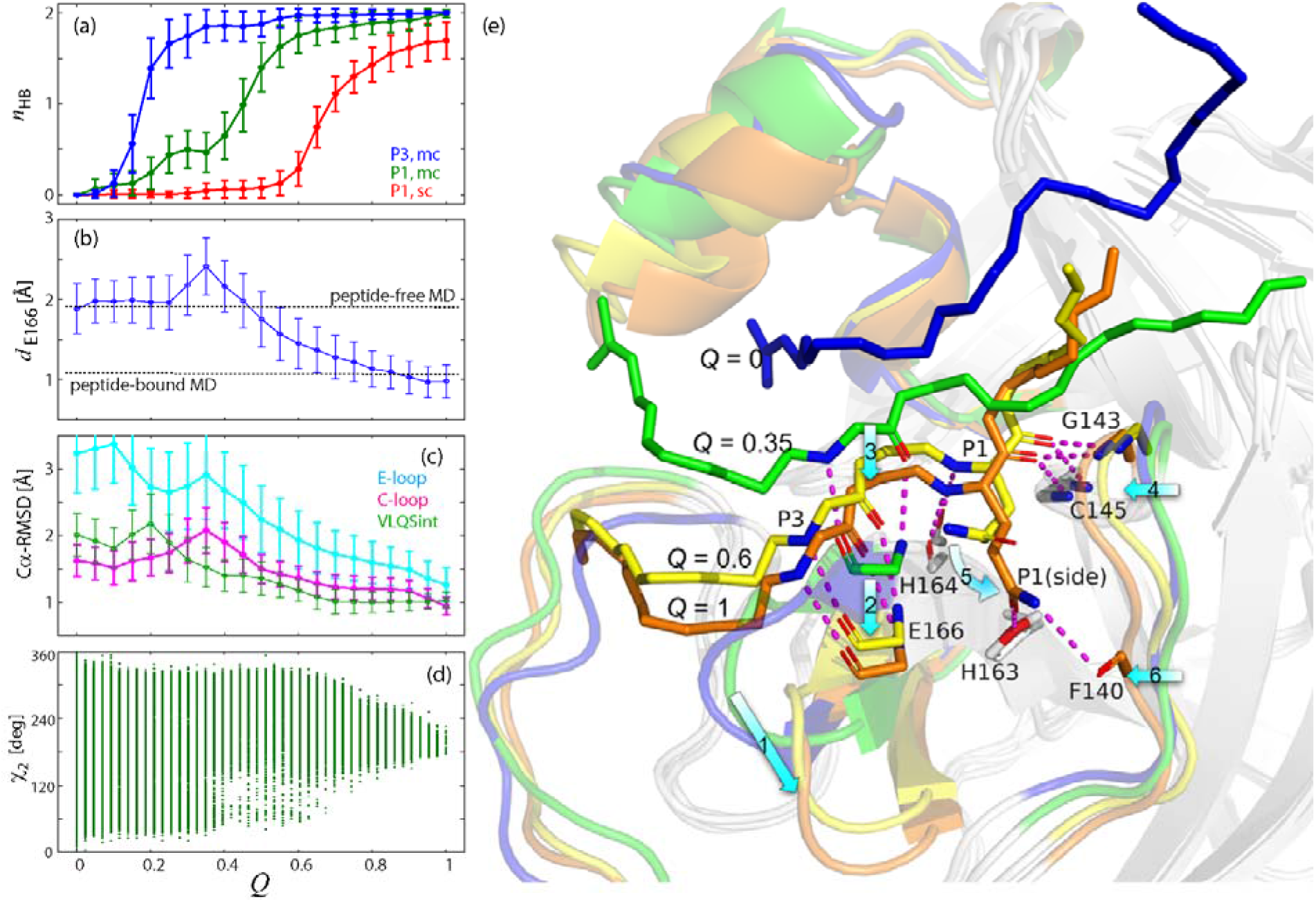
Peptide recognition process along native contact formations. (a) The number of hydrogen-bond *n_HB_* as a function of *Q* for group1 (the main-chain of P3_Val; HB #1and #2 in Table 1; blue), group2 (the main-chain of P1_Gln; #3 and #5; green) and group3 (the side-chain of P1_Gln; #6 and #7, red). (b) The distance of the Glu166 Cα atom from that of the peptide-bound state, *d*_E166_. (c) The Cα-RMSD values of the E-loop (cyan) and the C-loop (red) after the superimposition to the core region, and the Cα-RMSD of the peptide VLQS residues (green). (d) The sampled χ2 angle in P1_Gln indicated by dots. (e) Representative structures during the peptide binding at *Q* = 0 (blue), 0.35 (green), 0.6 (yellow) and 1 (orange). The main-chain of the peptide and side-chain of P1_Gln (Q = 0.6 and 1) are shown on the stick. The C-loop, E-loop and Linker are drawn by a coloured cartoon and the other part of the protein by a white cartoon. The HBs listed in Table 1 are indicated by magenta broken lines. The structural changes during the binding process are marked by cyan arrows with the numbers 1–6: Glu166 catches P3_Val at the outer region of the cleft (*Q* = 0.35) to form the HBs in group 1 (see Table 1), and the E-loop’s motion toward active form (Arrow 1) leads Glu166 (Arrow 2) and P3_Val (Arrow 3) to the binding site. This change accompanies the motion of P1_Gln toward the HB site of group 2 (including the oxyanion hole) causing the shift of the C-loop (Arrow 4; *Q* = 0.6). Finally, the fluctuation of the P1_Gln side-chain is converged to the peptide-bound form and the formation of the HBs in group 3 are complete (Arrow 5; *Q* = 1), which accompany the motion of the C-loop at Phe140 (Arrow 6).

Based on the observation in Fig. 4a, the structural basis of the peptide recognition was further elucidated by relating the sequential formation of the HBs to the protein structural change and the peptide folding toward the peptide-bound state (Figs. 4b-d). A series of snapshots of the peptide binding process along *Q*-value (*Q* = 0, 0.35, 0.6 and 1) were employed to illustrate the derived findings (Fig. 4e).

At the initial stage of the HB formation at *Q* ~ 0.2, one of the HBs in the group 1 between Glu166 (main-chain) and P3_Val (main-chain) is firstly formed (Fig. 4a), where Glu166 is at the position of the peptide-free state (because the Cα distance of Glu166 from the peptide-bound state, *d*_E166_, was evaluated to be 1.9 ± 0.4 Å in the peptide-free MD; Fig. 4b). This HB triggers the peptide’s folding to form the peptide-bound structure, or initiates the decrease of the Cα-RMSD for the peptide VLQS residues from ~2 Å (Fig. 4c).

Glu166 is then more attracted to the peptide when the two HBs in the group 1 between Glu166 (main-chain) and P3_Val (main-chain) are complete at Q ~ 0.35 (Figs. 4b and 4e), accompanying the increase of *d*_E166_ (fig. 4d). This overshoot of Glu166 appears as if the E-loop goes out of the cleft to catch the peptide located outside the binding site. This dynamic behavior of the E-loop reminds us of the fly-casting mechanism (33), in which the unfolded and highly flexible region in a protein is used to bind a ligand to form the complex structure (though the Eloop does not unfold). Glu166 starts to cause a significant motion toward the peptide-bound state when *Q*-value becomes greater than 0.35 (Arrow 2 of Fig. 4e), seen in the decrease of *d*_E166_ to the level in the peptide-bound state (*d*_E166_ = 1.1 ± 0.3 Å calculated in the peptide-bound MD; Fig. 4b) and thus leads the peptide into the binding site buried inside of the cleft (Arrow 3 of Fig. 4e). These binding dynamics of Glu166 are mostly due to the E-loop’s structural change (residues 166–171) (Fig. 4c; Arrow 1 of Fig. 4e). Since in the peptide-free simulation the E-loop was highly flexible (see Fig. S2), the E-loop susceptibly responds to the interactions with the peptide to lead the peptide to the bound state. In our previous study, the E-loop was found to make a ligand-size dependent conformational change (17). This finding in the crystal structure ensemble corroborates the susceptible response to the peptide interactions observed in the WE simulation.

At *Q* ~ 0.6, the peptide motion led by the E-loop further causes the residues in the group 2 to form the HBs with P1_Gln (Figs. 4a and 4e). These HBs are formed during a slight shift of the C-loop at Gly143 and Cys145 toward the peptide (Arrow 4 of Fig. 4e), seen in the decrease of the C-loop Cα-RMSD (Fig. 4d). However, at *Q* ~ 0.6, the side-chain of P1_Gln still largely fluctuates (Fig. 4d) not allowing to fix it at the binding conformation.

Finally, at *Q* ~ 1, the final stage of the binding process, the conformational fluctuation of the P1_Gln side-chain is gradually converged to the peptide-bound form (Fig. 4d) and the HBs in the group 3 are formed to complete the binding process (Arrow 5 of Fig. 4e). The binding of the side-chain of P1_Gln with Phe140 accompanies a further shift of the C-loop at Phe140 (Arrow 6 of Fig. 4e).

## Discussion

The path sampling simulations of the eight-residue peptide binding/unbinding process to/from SARS-CoV-2 3CL^pro^ were conducted using the WE method (24–26). The binding kinetics was successfully quantified in terms of the MFPT. The obtained MFPT showed reversible kinetics in the peptide binding/unbinding process except for the free energy difference favouring the peptide-bound state. It also underwent an extensive slowdown relative to simple diffusion along the reaction coordinate, RMSD_VLQS_. Thus, the kinetics was categorised as subdiffusion, which is supposed to be one of the common behavior of flexible peptides (32).

The path ensemble obtained in the WE simulations enabled us to explain the details of the peptide binding/unbinding process. The initial stage of the binding process starts with the diffusion on the protein surface that allows the overall rotation and a considerable conformational change of the peptide. The diffusion is directed toward the cleft region between the two domains in the Chymotrypsin fold to increase the number of non-specific contacts. This cleft plays a crucial role in leading the highly flexible peptide to the peptide-bound form: aligning the peptide to the cleft to restrain the overall rotation, increasing the number of contacts and enhancing the peptide’s desolvation.

After the peptide enters the cleft, the specific native contacts start to be formed first at the main-chain of Glu166, where the E-loop’s dynamics play a critical role. The E-loop is intrinsically flexible and shows a large conformational change upon ligand binding. The E-loop responds to the approaching peptide and causes a large shift toward the peptide to form the HBs of Glu166 (group 1 in Table 1), like the fly-casting mechanism (33). These HBs trigger the sequential formation of the HBs to P1_Gln main-chain (group2) to lead the peptide more deeply into the cleft. The last event to complete the binding process occurs in the side-chain of P1_Gln (the HBs in group 3). This is because the side-chain flexibility can be constrained only after the peptide’s residues and the protein surrounding the side-chain is fixed by the native contacts. During the binding process, first the cleft of the Chymotrypsin fold and then the specific native contacts gradually restrain the flexible peptide conformation to form the peptide-bound state. Thus, it is concluded that the peptide binding is the process of gradual damping of the motions of both the peptide and protein that is highly regulated by the non-specific and specific peptide-protein interactions.

## Methods

### Model construction and MD simulations

Conventional MD simulations were performed for the SARS-CoV-2 3CL^pro^ dimer with and without the bound peptide to study the dynamics at the termini of the path ensemble calculated by the subsequent WE simulation. The simulation model was taken from the SARS-CoV crystal structure (PDB: 2q6g; 34), which binds the peptide substrate (residues 3258–3265 of P0DTC1 or P0DTD1, TSAVLQ↓SG capped with the terminal acetyl and N-methyl groups (Mw = 816.9); the C-terminal 6 residues of nsp4 and the N-terminal 2 residues of nsp5 (3CL^pro^) in the polyprotein of SARS-CoV-2; ‘↓’ indicates the substrate cleavage site and thus the peptide corresponds to P6-P1↓P1 ‘P2’; the three C-terminal residues in 2q6g were excluded from the model). The 12 SARS-CoV-specific amino acids with the mutated Ala41 in 2q6g were converted into those of SARS-CoV-2 employing MODELLER (35). This model was selected because it has a unique structure representing the non-covalent interaction between Cys145 and the substrate (17,34). The other complex crystal structures with non-covalent interactions with the substrate peptide (PDB:1z1j, 5b6o, 6xoa and 7joy) have mutations, C145A or C145S. Recently, the structure of the 3CL^pro^ (H41A) complexed with the same peptide was released (PDB:7dvp;12). The structure of 7dvp was discovered to be very similar to that of 2q6g; Cα-RMSD (residues 1–200) = 0.38 Å and RMSD (non-hydrogen atoms of the eight residues of the peptide after superposition of the protein) = 0.70 Å.

Rectangular simulation boxes were constructed with a margin of 12 Å to the boundary of the simulation box and fully solvated by 25,955 TIP3P water molecules (36) and eight sodium ions to neutralise the simulation systems, resulting in 89,501 atoms. The molecular recognition and the proteolytic reactions occur seamlessly, and the substrate peptide forms a covalently bound intermediate between Cys145 Sγ of 3CL^pro^ and P1_Gln O during the proteolysis reaction (14). However, due to the limitation of the MD simulation, 3CL^pro^ and the peptide interact with each other in a non-covalent manner throughout the simulations. The peptide-unbinding simulation was conducted with the same system after removing the peptides.

AMBER ff14SB (37) was employed for the potential energy of the protein. The MD simulations were conducted using AMBER 16 (38) under the constant temperature and pressure (NPT) condition at *P* = 1 atm and *T* = 300 K, using Berendsen’s barostat and Langevin dynamics to control the temperature with 1.0 ps^−1^ as the collision frequency. The particle mesh Ewald method (39) was used for the electrostatic interactions. The time step was 2 fs with the constraints of bonds involving hydrogen atoms via the SHAKE algorithm (40). For each of the peptide-bound and peptide-free systems, the simulation was performed for 1 μs and three times starting with different initial velocities. The trajectories were recorded at 10 ps intervals.

### Construction of Motion Tree

In the analysis of the trajectories derived in the MD simulations of the peptide-bound and peptide-free states, protein dynamics were analyzed as a set of collective motions among residue clusters behaving as rigid structural units. These clusters were defined by the hierarchical clustering of the fluctuations of inter-residue distances to construct a dendrogram named *‘Motion Tree’* (27,28). Each node of the Motion Tree shows a pair of clusters mutually fluctuating with the magnitude represented by the node height. In previous studies, the Motion Tree was applied to the analyses of the trajectories of MD simulations (28,41,42) and crystal structure ensemble (17,18).

The metric of the hierarchical clustering for the Motion Tree comparing two structures (snapshots near the average structures of the peptide-bound and peptide-free simulations) was defined by *D_mn_* = |*d_mn_*^bound^ – *d_mn_*^free^|, where *d_mn_* represents the distance between the Cα atoms of residues *m* and *n* (27). Structural fluctuations of the peptide-bound structure or the peptide-free structure during the simulation were described by employing the metric of 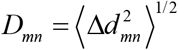, where 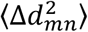 represents the variance of the structural ensemble obtained in the simulation (28). Since 3CL^pro^ forms a homodimer, *D_mn_* was symmetrized by averaging the duplicated structures of AB and BA, where AB represents the original dimer of protomers A and B, and BA is the exchanged dimer. The C-terminal segment (residues 301–306) was excluded in the analysis since it is highly flexible and irrelevant to 3CL^pro^ function.

### WE simulations

The path ensembles of the binding and unbinding processes were obtained using the WE simulation (26). The WE simulation carries out a number of short-time unbiased MD runs repeatedly, whose trajectories are then connected to construct full continuous paths progressing along a pre-defined reaction coordinate. The original Huber-Kim algorithm (24,26) of the WE simulation was employed for the binding/unbinding simulation to/from protomer A, while the peptide on protomer B was kept in the peptide-bound form (see Fig. 1a). It was confirmed in the preliminary simulations that there was no detectable cooperativity between the two binding sites. As the one-dimensional reaction coordinate in the WE simulation, we chose the root-meansquare deviation (RMSD) of the non-hydrogen atoms of the substrate residues VLQS (corresponding to P3-P1’) from the crystal structure after superimposing the protein’s core region (17; hereinafter designated as RMSD_VLQS_). Thus, RMSD_VLQS_ reflects not only the peptide internal conformation but also the peptide’s overall translation and rotation. The discretization of the reaction coordinate for the WE simulation was conducted as follows: RMSD_VLQS_ < 2 Å is defined as the peptide-bound state, RMSD_VLQS_ > 25 Å is the peptide-free state, and 40 bins are set between 2 and 12 Å with the interval of 0.25 Å and 26 bins between 12 and 25 Å with the interval of 0.5 Å; the number of bins, *M*_bin_, is thus 68. The peptide-bound state corresponds to the range of thermal fluctuation seen in the MD simulation of the peptide-bound form, while the peptide-free form is the peptide position sufficiently distant from the binding site with no interaction with the protein. As the initial condition for the simulation, the configuration was set at bin 1 (the peptide-free state for the binding simulation and the peptide-bound state for the unbinding simulation), i.e. *ρ*_1_(0) = 1, where *ρ_I_*(*t*) represents the probability of the occupancy at bin *I* (*I* = 1, 2,..*M*_bin_) at time *t*. To evolve *ρ_I_*(*t*), 16 unbiased MD simulations of *τ* (= 25 ps in this study) were performed starting from a representative set of 16 copies of the system involved in bin *I* when *ρ_I_*(*t*) > 0. Thus, at time *t*, 1088 (= 16 copies × 68 bins) MD simulations were conducted. This procedure was repeated *L* times (= 300 for the binding simulation and 400 for the unbinding simulation) iteratively to evolve *ρ_I_*(*t*), i.e. the time interval of *ρ_I_*(*t*) is thus *L_τ_* = 7.5 ns and 10 ns for binding and unbinding simulations, respectively. In this research, the unbinding paths were first computed by starting at a structure taken randomly from the trajectories of the MD simulation of the peptide-bound form and then the computation of the binding paths was performed starting from each of the three separate structures of the peptide-free form generated by the unbinding WE runs (see Fig. S3). The WE runs were repeated three times for the error estimation of the rate constants. In total, the simulation time amounts to 24.5 μs (25 ps × 16 copies × 68 bins × 300 iterations × 3 runs) for the binding simulation and 32.6 μs (25 ps × 16 copies × 68 bins × 400 iterations × 3 runs) for the unbinding simulation.

The MFPT, or the inverse of the rate constant *k*, was calculated from the probability flow, *p*_flow_, as follows:

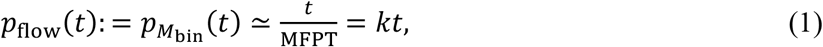

where the approximation is for a sufficiently small *t*. First, *ρ_I_*(*t*) was time-averaged with a timestep of 100 ps to reduce fluctuating noise. Then, the averaged *ρ_I_*(*t*) at the terminal bin (*I* = *M*_bin_), corresponding to *p*_flow_, was employed for the calculation of gradient by linear regression in the range of 5 ≤ *t* ≤ 7.5 ns for the binding simulation and 7.5 ≤ *t* ≤ 10 ns for the unbinding simulation (see Fig. 2a). The binding rate constant *k*_off_ and *k*_on_ were calculated respectively, using

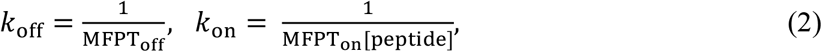

where MFPT_on_ and MFPT_off_ represent the values of MFPT obtained in the binding and unbinding simulations, respectively and [peptide] is the peptide’s concentration in the simulation box, [peptide] = (1 molecule)/(the volume of the simulation box) = 1.9 × 10^−3^ M. The peptide-bound state in the computation of the MFPT was redefined according to the HBs based on Table 1, instead of the RMSD_VLQS_ value, i.e. by using the conditions that the HBs #3 and #5 in the group 2 and more than one HB (#6 or #7) in the group 3 are formed.

## Data availability

The data that support the findings of this study are available from the corresponding author upon reasonable request.

## Acknowledgements

This research was supported by Platform Project for Supporting Drug Discovery and Life Science Research (Basis for Supporting Innovative Drug Discovery and Life Science Research (BINDS)) from AMED under Grant Number JP21am0101109, and by Research Support Project for Life Science and Drug Discovery (BINDS) from AMED under Grant Number JP22ama121023. K.M. was supported by Grant for Basic Science Research Projects from The Sumitomo Foundation (190636), and by MEXT Grants-in-Aid for Scientific Research (C), 20K03883. The computations were performed at Graduate School of Medical Life Science at Yokohama City University.

## Author contributions

K.M., T.E., M.I. and A.K. designed research; K.M. performed research; K.M. and A.K. analyzed data; K.M., T.E., M.I. and A.K. wrote the paper.

## Competing interests

The authors declare that they have no conflict of interest.

## Additional information

Supplementary information is available for this paper at …

## Notes

### Competing Interest Statement

The authors have declared no competing interest.

### Summary of Updates

The manuscript has been proofread by a English editing company, after substantial modifications were added.

## References

1. Rawlings ND, Barrett AJ, Thomas PD, Huang X, Bateman A, Finn RD. The MEROPS database of proteolytic enzymes, their substrates and inhibitors in 2017 and a comparison with peptidases in the PANTHER database. Nucleic Acids Res 2018;46:D624–D632.

2. Ullrich S, Nitsche C. The SARS-CoV-2 main protease as drug target. Bioorg Med Chem Lett 2020;30:127377.

3. Banerjee R, Perera L, Tillekeratne LMV. Potential SARS-CoV-2 main protease inhibitors. Drug Discov Today 2021;26:804–816.

4. Vandyck K, Deval J. Considerations for the discovery and development of 3-chymotrypsin-like cysteine protease inhibitors targeting SARS-CoV-2 infection. Curr Opin Virol 2021;49:36–40.

5. Tan K, Maltseva NI, Welk LF, Jedrzejczak RP, Joachimiak A. The crystal structure of 3CL MainPro of SARS-CoV-2 with C145S mutation, http://www.rcsb.org/structure/6XOA; 2020.

6. Lee J, Worrall LJ, Vuckovic M, Rosell FI, Gentile F, Ton AT, Caveney NA, Ban F, Cherkasov A, Paetzel M, Strynadka NCJ. Crystallographic structure of wild-type SARS-CoV-2 main protease acyl-enzyme intermediate with physiological C-terminal autoprocessing site. Nat Commun 2020;11:5877.

7. Ferre RA, Liu W, Ryan K, Grodsky N, Gajiwala KS. SARS coronavirus-2 main protease dimer auto-processes N-terminus in cis and C-terminus in trans, http://www.rcsb.org/structure/7LMC; 2021.

8. MacDonald EA, Frey G, Namchuk MN, Harrison SC, Hinshaw SM, Windsor IW. Recognition of divergent viral substrates by the SARS-CoV-2 main protease. ACS Infect Dis 2021;7:2591–2595.

9. Noske GD, Nakamura AM, Gawriljuk VO, Fernandes RS, Lima GMA, Rosa HVD, Pereira HD, Zeri ACM, Nascimento AFZ, Freire MCLC, Fearon D, Douangamath A, von Delft F, Oliva G, Godoy AS. A Crystallographic snapshot of SARS-CoV-2 main protease maturation process. J Mol Biol 2021;433:167118.

10. Kneller DW, Zhang Q, Coates L, Louis JM, Kovalevsky A. Michaelis-like complex of SARS-CoV-2 main protease visualized by room-temperature X-ray crystallography. IUCrJ 2021;8:973–979.

11. Noske GD, Song Y, Fernandes RS, Oliva G, Godoy AS. Cryo-EM structure of SARS-CoV-2 Main protease C145S in complex with N-terminal peptide, http://www.rcsb.org/structure/7S82; 2022.

12. Zhao Y, Zhu Y, Liu X, Jin Z, Duan Y, Zhang Q, Wu C, Feng L, Du X, Zhao J, Shao M, Zhang B, Yang X, Wu L, Ji X, Guddat LW, Yang K, Rao Z, Yang H. Structural basis for replicase polyprotein cleavage and substrate specificity of main protease from SARS-CoV-2. Proc Natl Acad Sci U S A 2022;119:e2117142119.

13. Schechter I, Berger A. On the size of the active site in proteases. I. Papain. Biochem Biophys Res Commun 1967;27:157–162.

14. Otto HH, Schirmeister T. Cysteine proteases and their inhibitors. Chem Rev 1997;97:133–172.

15. Vankadara S, Wong YX, Liu B, See YY, Tan LH, Tan QW, Wang G, Karuna R, Guo X, Tan ST, Fong JY, Joy J, Chia CSB. A head-to-head comparison of the inhibitory activities of 15 peptidomimetic SARS-CoV-2 3CLpro inhibitors. Bioorg Med Chem Lett 2021;48:128263.

16. Owen DR, Allerton CMN, Anderson AS, Aschenbrenner L, Avery M, Berritt S, Boras B, Cardin RD, Carlo A, Coffman KJ, Dantonio A, Di L, Eng H, Ferre R, Gajiwala KS, Gibson SA, Greasley SE, Hurst BL, Kadar EP, Kalgutkar AS, Lee JC, Lee J, Liu W, Mason SW, Noell S, Novak JJ, Obach RS, Ogilvie K, Patel NC, Pettersson M, Rai DK, Reese MR, Sammons MF, Sathish JG, Singh RSP, Steppan CM, Stewart AE, Tuttle JB, Updyke L, Verhoest PR, Wei L, Yang Q, Zhu Y. An oral SARS-CoV-2 Mpro inhibitor clinical candidate for the treatment of COVID-19. Science 2021;374:1586–1593.

17. Kidera A, Moritsugu K, Ekimoto T, Ikeguchi M. Allosteric regulation of 3CL protease of SARS-CoV-2 and SARS-CoV observed in the crystal structure ensemble. J Mol Biol 2021;433:167324.

18. Moritsugu K, Nishino Y, Kidera A. Inter-lobe motions allosterically regulate the structure and function of EGFR kinase. J Mol Biol 2020;432:4561–4575.

19. Renaud JP, Chung CW, Danielson UH, Egner U, Hennig M, Hubbard RE, Nar H. Biophysics in drug discovery: Impact, challenges and opportunities. Nat Rev Drug Discov 2016;15:679–698.

20. Paul F, Wehmeyer C, Abualrous ET, Wu H, Crabtree MD, Schöneberg J, Clarke J, Freund C, Weikl TR, Noé F. Protein-peptide association kinetics beyond the seconds timescale from atomistic simulations. Nat Commun 2017;8:1095.

21. Nunes-Alves A, Kokh DB, Wade RC. Recent progress in molecular simulation methods for drug binding kinetics. Curr Opin Struct Biol 2020;64:126–133.

22. Copeland RA, Pompliano DL, Meek TD. Drug target residence time and its implications for lead optimizationzation. Nat Rev Drug Discov 2006;5:730–739.

23. Copeland RA. The drug-target residence time model: A 10-year retrospective. Nat Rev Drug Discov 2016;15:87–95.

24. Huber GA, Kim S. Weighted-ensembleeighted-ensemble Brownian dynamics simulations for protein association reactions. Biophys J 1996;70:97–110.

25. Zwier MC, Pratt AJ, Adelman JL, Kaus JW, Zuckerman DM, Chong LT. Efficient atomistic simulation of pathways and calculation of rate constants for a protein-peptide binding process: Application to the MDM2 protein and an intrinsically disordered p53 peptide. J Phys Chem Lett 2016;7:3440–3445.

26. Zuckerman DM, Chong LT. Weighted ensemble simulation: Review of methodology, applications, and software. Annu Rev Biophys 2017;46:43–57.

27. Koike R, Ota M, Kidera A. Hierarchical description and extensive classification of protein structural changes by Motion Tree. J Mol Biol 2014;426:752–762.

28. Moritsugu K, Koike R, Yamada K, Kato H, Kidera A. Motion Tree delineates hierarchical structure of protein dynamics observed in molecular dynamics simulation. PLOS ONE 2015;10:e0131583.

29. Ma S, Devi-Kesavan LS, Gao J. Molecular dynamics simulations of the catalytic pathway of a cysteine protease: A combined QM/MM study of human cathepsin K. J Am Chem Soc 2007;129:13633–13645.

30. Kneller DW, Phillips G, Weiss KL, Pant S, Zhang Q, O’Neill HM, Coates L, Kovalevsky A. Unusual zwitterionic catalytic site of SARS-CoV-2 main protease revealed by neutron crystallography. J Biol Chem 2020;295:17365–17373.

31. Bollt EM, ben-Avraham D. What is special about diffusion on scale-free nets? New J Phys 2005;7:26.

32. Neusius T, Daidone I, Sokolov IM, Smith JC. Subdiffusion in peptides originates from the fractal-like structure of configuration space. Phys Rev Lett 2008;100:188103.

33. Shoemaker BA, Portman JJ, Wolynes PG. Speeding molecular recognition by using the folding funnel: The fly-casting mechanism. Proc Natl Acad Sci U S A 2000;97:8868–8873.

34. Xue X, Yu H, Yang H, Xue F, Wu Z, Shen W, Li J, Zhou Z, Ding Y, Zhao Q, Zhang XC, Liao M, Bartlam M, Rao Z. Structures of two coronavirus main proteases: Implications for substrate binding and antiviral drug design. J Virol 2008;82:2515–2527.

35. Fiser A, Sali A. MODELLER: Generation and refinement of homology-based protein structure models. Method Enzymol 2003;374:461–491.

36. Jorgensen WL, Chandrasekhar J, Madura JD, Impey RW, Klein ML. Comparison of simple potential functions for simulating liquid water. J Chem Phys 1983;79:926–935.

37. Maier JA, Martinez C, Kasavajhala K, Wickstrom L, Hauser KE, Simmerling C. ff14SB: Improving the accuracy of protein side chain and backbone parameters from ff99SB. J Chem Theor Comput 2015;11:3696–3713.

38. Case DA, Cheatham TE, Darden T, Gohlke H, Luo R, Merz KM, Onufriev A, Simmerling C, Wang B, Woods RJ. The Amber biomolecular simulation programs. J Comput Chem 2005;26:1668–1688.

39. Darden T, York D, Pedersen L. Particle mesh Ewald – An Nlog(N) method for Ewald sums in large systems. J Chem Phys 1993;98:10089–10092.

40. Ryckaert JP, Ciccotti G, Berendsen HJC. Numerical integration of the Cartesian equations of motion of a system with constraints: Molecular dynamics of n-alkanes. J Comput Phys 1977;23:327–341.

41. Moritsugu K, Ito T, Kidera A. Allosteric response to ligand binding: Molecular dynamics study of the N-terminal domains in IP3 receptor. Biophys Physicobiol 2019;16:232–239.

42. Koike R, Takeda S, Maéda Y, Ota M. Comprehensive analysis of motions in molecular dynamics trajectories of the actin capping protein and its inhibitor complexes. Proteins 2016;84:948–956.

