## Supporting Information for "Binding and unbinding pathways of peptide substrate on SARS-CoV-2 3CL protease"

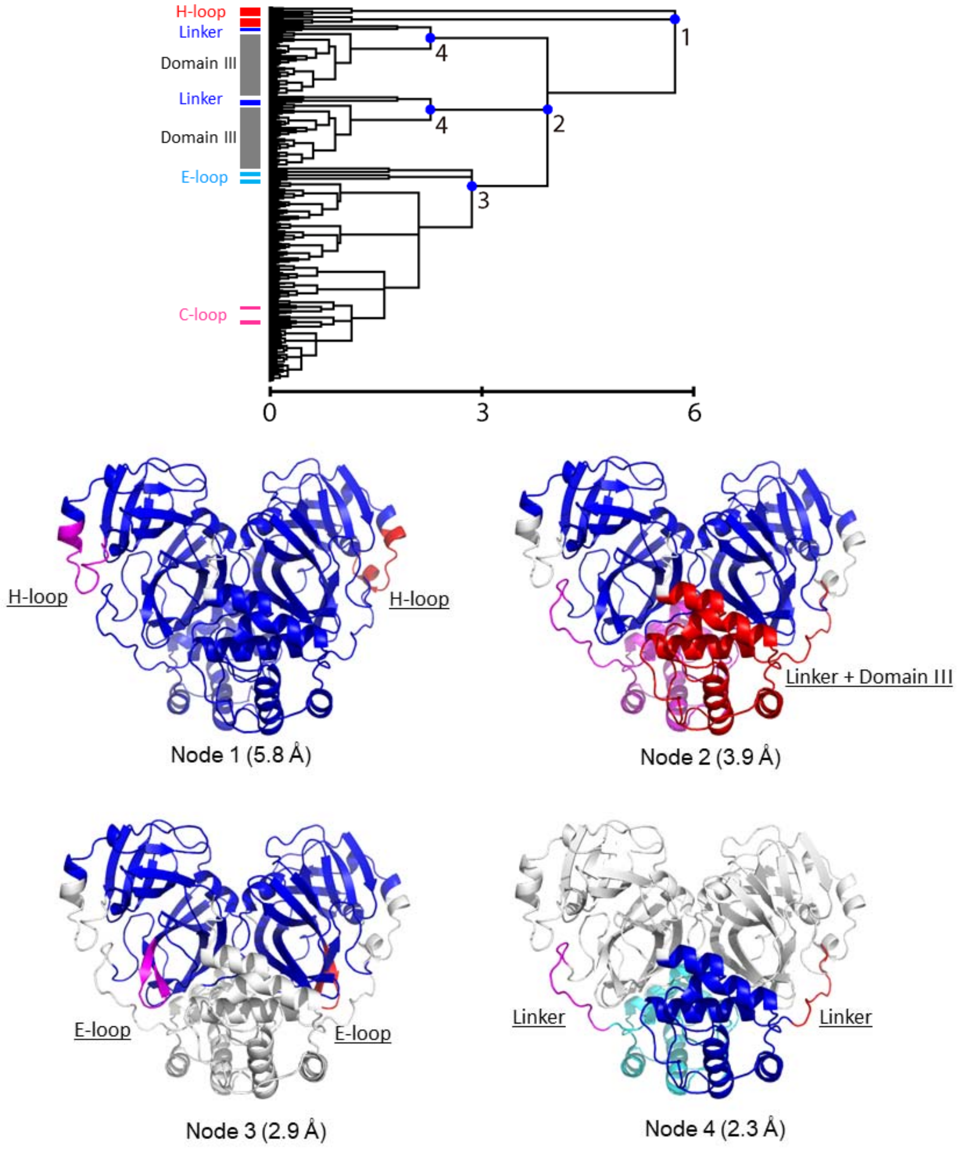


**Figure S1.** Motion Tree describing the difference between the two structures near each average structure taken from the MD simulations of the peptide-bound and peptide-free 3CL$^{pro}$ dimers (27). The positions of the moving clusters are labelled at the end of the tree. A branching node of the Motion Tree consists of a larger part (blue/cyan) and a smaller part (red/magenta) changing their mutual position as rigid bodies with the amplitudes of the MT scores represented by the node height.

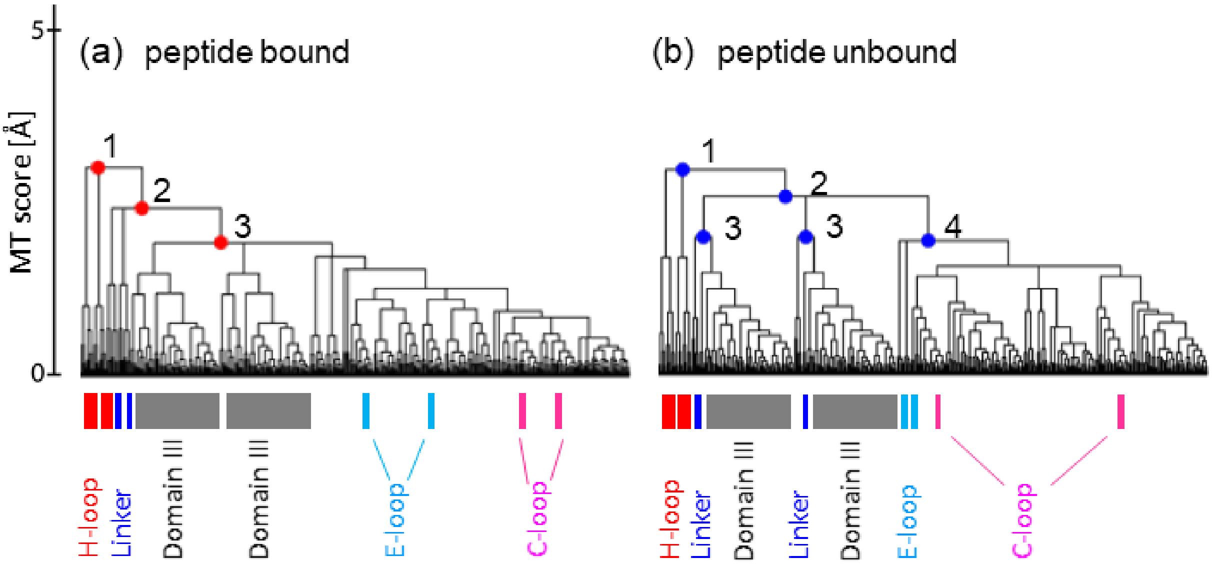


**Figure S2.** Motion Trees describing dynamic fluctuations of (a) the peptide-bound and (b) the peptide-free 3CL$^{pro}$ dimers, constructed from the variance of the inter-residue distances observed in the 3 × 1-μs MD simulations (28). Nodes of the tree represent the motions of the moving clusters relative to the protein core: H-loop, Linker, domain III, E-loop. The E-loop in the peptide-bound simulation and the C-loop in both simulations are within the rigid core region of domains I and II with small MT scores.

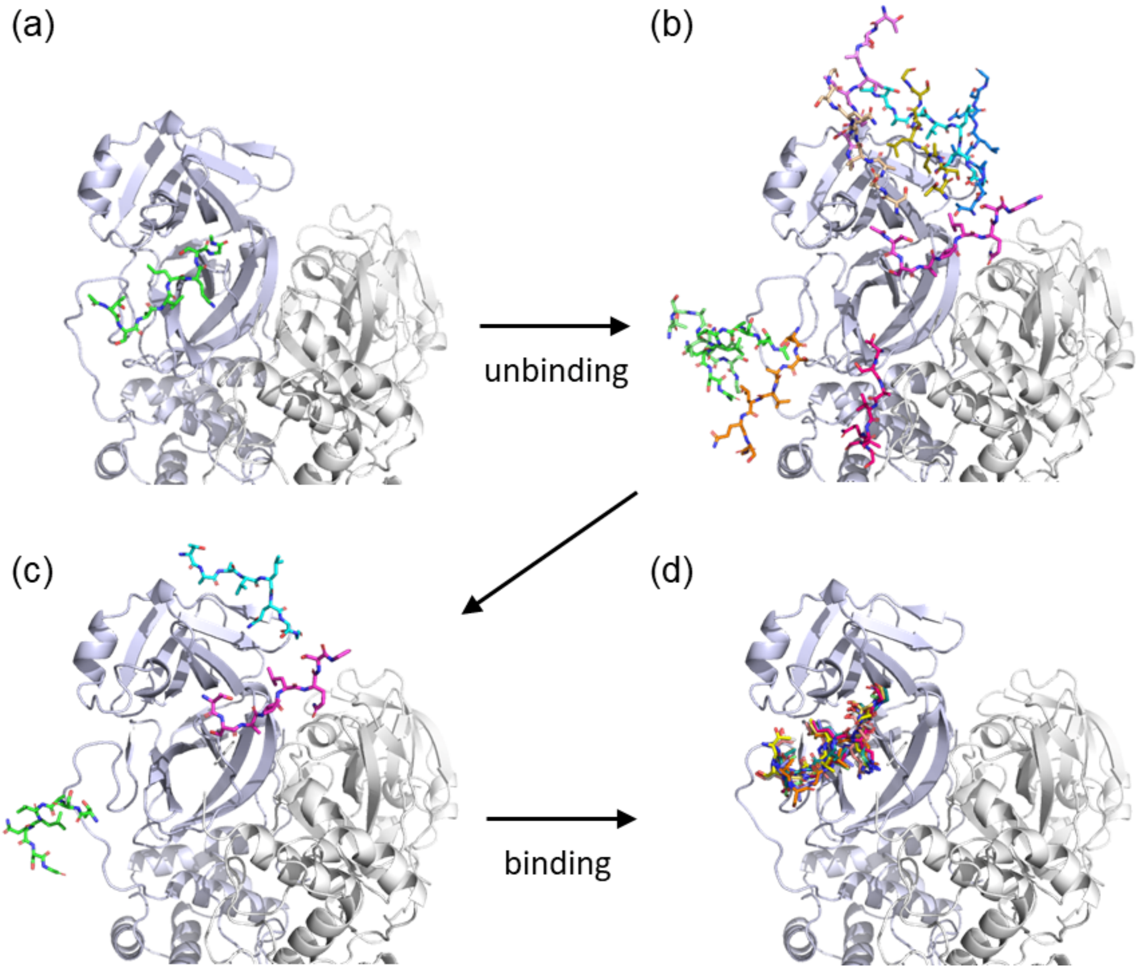


**Figure S3.** The WE simulation setup. (a) The unbinding WE runs were started from a peptide-bound form taken from the peptide-bound simulation. (b) Representative nine structures in the terminal bin show widely distributed unbinding paths. (c) The binding WE runs were started from three peptide positions taken randomly from the terminal bin of the unbinding WE runs. (d) Representative nine structures in the terminal bin show the convergence of the binding paths to the peptide-bound form.

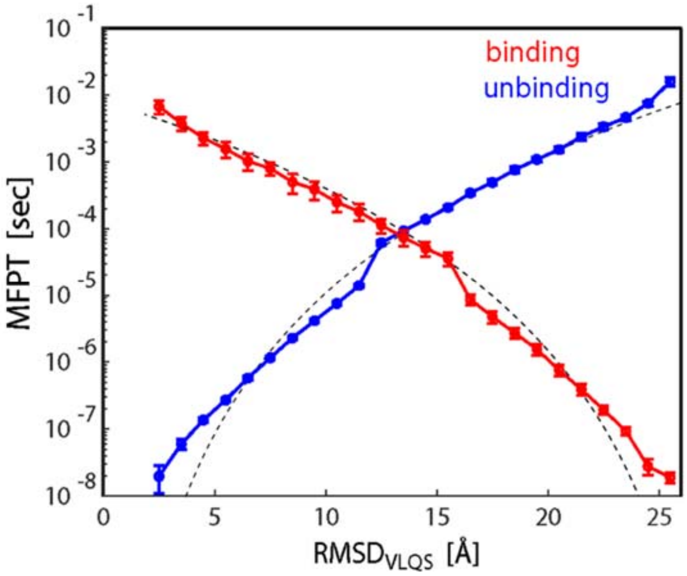


**Figure S4**. MFPT values for the binding and unbinding processes along the reaction coordinate, RMSD$_{VLQS}$ (= $r$). The broken curves are $r^d$ for the unbinding process and $(r_{max}-r)^d$ for the binding process with $d$ = 7. See text for the definition.

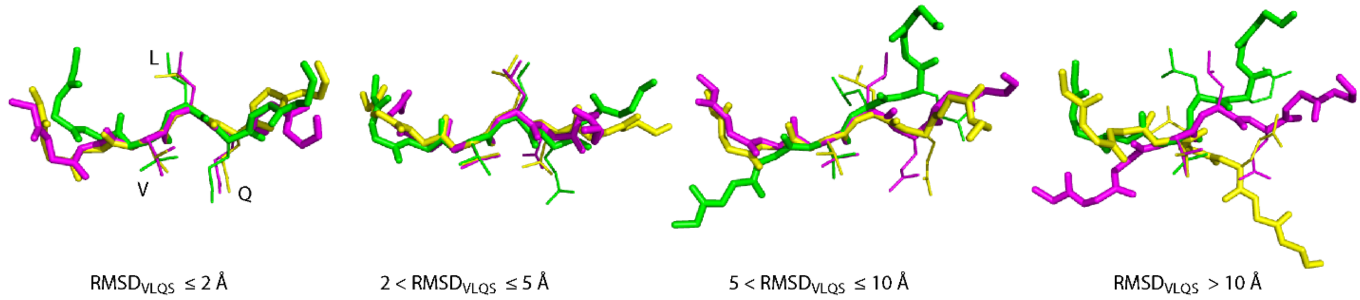


**Figure S5.** Representative peptide structures obtained from the WE simulations are shown separately in the designated RMSD$_{VLQS}$ ranges. These structures are superimposed at the Cα atoms of the residues VLQS.

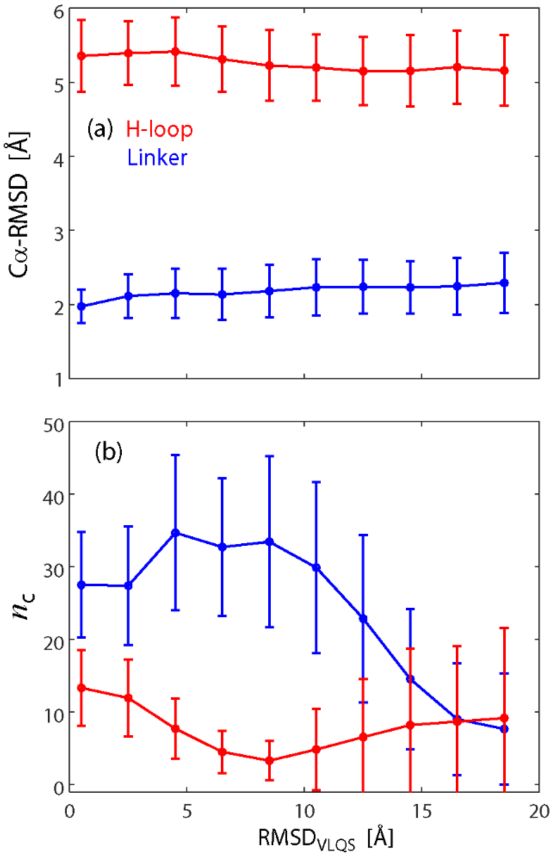


**Figure S6.** (a) Cα-RMSD and (b) the numbers of the atom contacts with the peptide, $n_C$, are plotted against RMSD$_{VLQS}$; the H-loop (red) and the Linker (blue).

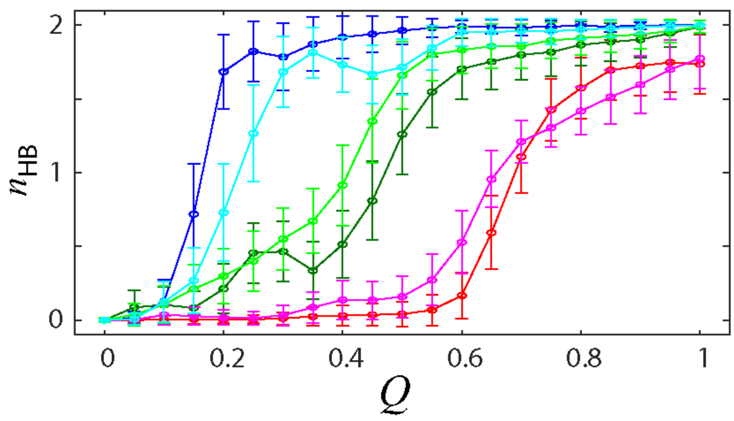


**Figure S7.** The numbers of the HBs, $n_{HB}$, for group 1 (cyan/blue), group 2 (light green/green) and group3 (magenta/red) during the binding/unbinding process. The binding process demonstrates higher Q values for group 1, while for groups 2 and 3, the unbinding process shows higher Q values, demonstrating that the two processes are reversible within the error range.

**Table S1. Probability of hydrogen-bond formation $p_{HB}$ with C-loop during the MD simulations.**

| C-loop | Partner | $p_{HB,bound}$[a] | $p_{HB,free}$[b] |
|---|---|---|---|
| Gly138N | His172N$_{\delta 2}$ | 0.71 | 0.64 |
| Ser139N | Tyr126OH | 0.68 | 0.85 |
| Phe140O | Ser1(B)N[c] | 0.75 | 0.69 |
| Leu141N | Tyr118OH | 0.87 | 0.90 |
| Gly143O | Asn28N$_{\delta 2}$ | 0.70 | 0.63 |

[a] $p_{HB,bound}$ for the probability of occurrence of HBs during the peptide-bound MD
[b] $p_{HB,free}$ for the probability of occurrence of HBs during the peptide-free MD.
[c] (B) indicates the residue in another protomer.

**Table S2. Dynamic correlations of the moving clusters during the MD simulations.**

| Moving cluster | Dynamic correlation* | | | | |
|---|---|---|---|---|---|
| | C-loop | E-loop | H-loop | Linker | Domain III |
| C-loop | | 0.21 | 0.06 | 0.09 | 0.09 |
| E-loop | 0.11 | | 0.04 | 0.21 | 0.07 |
| H-loop | 0.05 | 0.07 | | 0.07 | 0.06 |
| Linker | 0.09 | 0.13 | 0.13 | | 0.08 |
| domain III | 0.13 | 0.05 | 0.05 | 0.12 | |

* Dynamic correlation calculated as the average absolute values of the correlation coefficients of atomic displacement between a pair of moving clusters. The upper right entries are for the peptide-bound MD and the lower-left entries are for the peptide-free MD.

**Table S3. Kinetic parameters of the enzymatic reaction of SARS-CoV-2 3CL$^{pro}$ for the substrates whose sequences contain (TS)AVLQSG**

| $K_M$ [μM] | $k_{cat}$ [$s^{-1}$] | substrate | Ref. |
|---|---|---|---|
| 28.2 | 0.16* | Dabcyl-KTSAVLQSGFRKME-Edans | 1 |
| 170 | 0.27* | Dabcyl-KTSAVLQSGFRKME-Edans | 2 |
| 5.086 | 1.349 | $His_6$-MBP-TSAVLQSGFRKM-mEYFP | 3 |
| 17.2 | 0.51 | Mca–AVLQSGFRK-DnpK | 4 |
| 10.5 | 2.2 | Dabcyl-KTSAVLQSGFRKME-Edans | 5 |
| 16.4 | 28 | Dabcyl-KTSAVLQSGFRKME-Edans | 6 |
| 41 | 0.52 | Dabcyl-KTSAVLQSGFRKME-Edans | 7 |
| 110.3 | 5.1 | Dabcyl-KTSAVLQSGFRKME-Edans | 8 |
| 35.36 | 0.39 | Dabcyl-KTSAVLQSGFRKME-Edans | 9 |
| 26.7 | 0.97 | Dabcyl-KTSAVLQSGFRKME-Edans | 10 |

* The value of $k_{cat}$ was calculated from $k_{cat}/K_M$.

**References**

1. Ma C, Sacco MD, Hurst B, et al. Cell Res. 2020;30:678-692
2. Kneller DW, Galanie S, Phillips G, *et al.* Structure. 2020;28:1313-1320.e3.
3. Miczi M, Golda M, Kunkli B, *et al.* Int J Mol Sci. 2020;21:9523.
4. Tripathi PK, Upadhyay S, Singh M, *et al.* Int J Biol Macromol. 2020;164:2622-2631
5. Kuo CJ, Chao TL, Kao HC, *et al.* Antimicrob Agents Chemother. 2021;65:e02577-20.
6. Noske GD, Nakamura AM, Gawriljuk VO, et al. J Mol Biol. 2021;433:167118.
7. MacDonald EA, Frey G, Namchuk MN, *et al.* ACS Infect Dis. 2021;7:2591-2595.
8. Abe, K, Kabe, Y, Uchiyama, S, *et al.* Sci Rep 2022;12:1299.
9. Sacco, MD, Hu, Y, Gongora, MV, *et al.* Cell Res 2022;32:498–500.
10. Greasley, SE, Noell, S, Plotnikova, O, *et al.* J. Biol. Chem. 2022:101972.
